# MGATRx: Discovering Drug Repositioning Candidates Using Multi-view Graph Attention

**DOI:** 10.1101/2020.06.29.171876

**Authors:** Jaswanth K. Yella, Anil G. Jegga

**Affiliations:** Department of Computer Science, University of Cincinnati; Division of Biomedical, Informatics, Cincinnati Children’s Hospital Medical, Center; Cincinnati OH USA; Division of Biomedical Informatics, Cincinnati Children’s Hospital Medical Center; Departments of Pediatrics and Computer Science, University of Cincinnati; Cincinnati OH USA

**Keywords:** Drug repositioning, Multi-view graphs, Graph neural network, Heterogeneous network

## Abstract

In-silico drug repositioning or predicting new indications for approved or late-stage clinical trial drugs is a resourceful and time-efficient strategy in drug discovery. However, inferring novel candidate drugs for a disease is challenging, given the heterogeneity and sparseness of the underlying biological entities and their relationships (e.g., disease/drug annotations). By integrating drug-centric and disease-centric annotations as multiviews, we propose a multi-view graph attention network for indication discovery (MGATRx). Unlike most current similarity-based methods, we employ graph attention network on the heterogeneous drug and disease data to learn the representation of nodes and identify associations. MGATRx outperformed four other state-of-art methods used for computational drug repositioning. Further, several of our predicted novel indications are either currently investigated or are supported by literature evidence, demonstrating the overall translational utility of MGATRx.

**CCS CONCEPTS:**

- Applied computing → Biological networks; Bioinformatics
- Computing methodologies → Semantic networks

**ACM Reference format:** Jaswanth K Yella, Anil G Jegga 2020. MGATRx: Discovering Drug Repositioning Candidates Using Multi-view Graph. In *Proceedings of BIOKDD’20: International Workshop on Data Mining for Bioinformatics.* San Diego, CA, USA, 9 pages.

## 1 Introduction

As per an estimate, the total average cost for developing a new drug – from discovery to approval - ranges from $2 billion - $3 billion and takes about 13–15 years [1]. To by-pass this conventional slow and cost-intensive procedure, computational approaches are increasingly used to identify new therapeutic uses (“drug repurposing” or “drug repositioning”) for existing or approved, late-stage clinical trial drugs or drugs that have failed for reasons other than safety. The premise is that drugs approved by the FDA or those in late stages of clinical trials have known safety and toxicity profiles and hence candidate repositioning drugs can enter clinical phases more rapidly and at a relatively reduced cost. Knowledge-based in-silico drug repurposing methods typically utilize drug and disease annotations to assess the similarity or connections between drugs and diseases as per of the novel indication discovery strategy.

Several network-based approaches have been developed for predicting novel drug-disease, disease-gene, and drug-target links with drugs, diseases or their annotations (genes, phenotypes, pathways, etc.) as nodes and their interaction or associations as edges. Recent developments in graph-based learning methods have been shown to improve prediction performance and have the potential to unveil hidden relationships between the nodes in heterogeneous graphs. For instance, several years ago Gottlieb et al. [2] proposed PREDICT which integrates drug-drug similarity (based on drug-protein sequence, and gene-ontology interactions) and disease-disease similarity (based on disease-phenotype and human phenotype ontology) to predict drug-disease associations. More recently, Himmelstein et al. [3] created hetionet a heterogeneous network integrating data from 29 public resources to identify drug repositioning candidates and predict the probability of treatment for drug-disease pairs [4]. Similarly, leveraging integrated drug annotation data, Liang et al. [5] used Laplacian regularized sparse subspace learning method. Zhang et al. [6] used similarity constrained matrix factorization method while Wang et al. [7] used neural networks for non-linear feature extraction to predict drug-disease association. Results from each of these studies show that integrating heterogeneous data from various sources can predict drug-disease associations while being robust to inherent noise in the networks used. Each of these similarity-based approaches was evaluated with an underlying assumption that nodes sharing similar annotations have similar associations. As a result, latent representations generated through similarity networks may not necessarily capture the complex relations of drugs and diseases. Additionally, similarity-based approaches do not consider missing information. Several drugs and diseases, for instance, have missing information and therefore are not considered. For example, drugs and diseases tend to have off-target or indirect effects either through protein interactions, signaling networks, or pathways which are not typically detected in the preclinical drug discovery stages.

To address the challenges related to missing information and information cascades, we adopt graph neural networks-based approach. While integrating multiple views will compensate for any missing information, it is essential for the algorithm to aggregate information from the node neighbors and *learn* latent robust representation. The graph convolutional network (GCN) proposed by Kipf et al. [8] showed superior results in generating node embeddings while simplifying convolutions in the spectral approach. The GCN performs localized first-order approximations of spectral graph convolutions which is primarily proposed for single-view homogeneous networks. More recently, MVGAE [9] and MedGCN [10] employed GCNs on multi-view networks for computational drug discovery. However, the quality of annotations varies across views that can potentially affect the aggregation mechanism. Biomedical and genomic data tend to be relatively sparse and therefore it is important to assign varying importance to different neighbors.

To address this issue, we further narrow down the aggregation mechanism in graph neural networks by using attention on multiview graphs. The graph attention proposed by [11], [12] has shown to be effective by selectively gathering relevant information from neighboring nodes. We extend this paradigm to multi-view graphs to learn meaningful and relevant information from their neighbors of different node types. Using the derived representation, we predict links between two nodes (i.e., drug and disease in this case) through multi-label classification. In this paradigm, without explicitly providing any additional features to each node-type, we leverage the importance of neighbor relevance in graph attention mechanism and label correlation information captured in a multi-label learning setting. The proposed framework and algorithm is built on curated sets of drugs (approved, investigational, and withdrawn), diseases, and their annotations compiled from various sources. For instance, excluding hetionet, most of the previous approaches for identifying novel drug-disease associations evaluated their performance using a dataset from Wang et. al [13], proposed in 2014. However, since 2014, several new drugs have been approved. In 2018, the FDA approved as many as 59 new drugs [14].

A systematic validation shows that the proposed algorithm, Multiview Graph Attention for drug repositioning (**MGATRx**), performs better than current state-of-the-art models. Furthermore, using a case study, we illustrate an insightful understanding of the drug mechanism of action by visualizing the attention weights of MGATRx.

## 2 Materials and Methods

### 2.1 Drug and Disease Annotation Datasets

#### Drugs

We initially collected small-molecules that are approved, investigational, and withdrawn from MedChemExpress (MCE) panel [15]. To improve the coverage, we also included drug annotations from the KEGG database. We normalized the drug names by mapping them to common identifiers to extract their annotations from other sources. We used DrugBank identifiers for this purpose as it also enabled us to capture additional drug annotations such as chemical structures, targets, and side-effects. We manually curated the compiled list and removed some classes of drugs (e.g., all topical applications) which we believe have relatively low repositioning potential. Our final compiled list of drugs included 4008 approved and investigational compounds and we used this list to extract their annotations from other sources (Table 1).

**Table 1:**
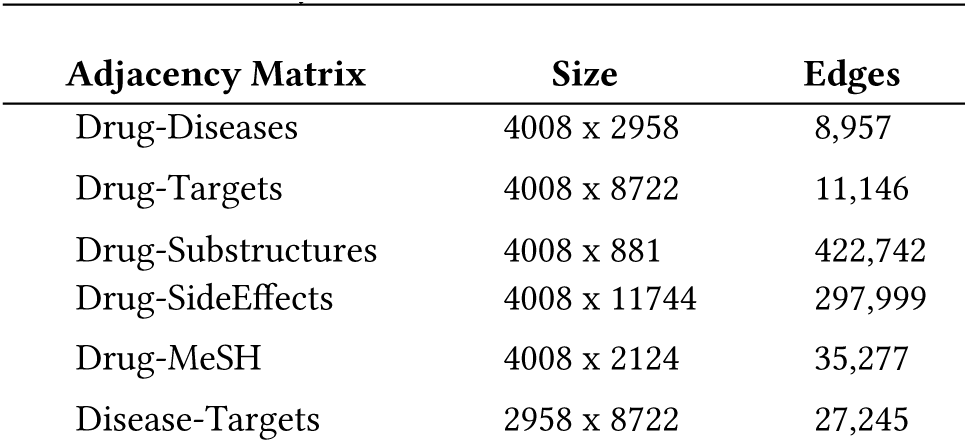
Graph information for each view type. Each adjacency matrix is considered as view and the size is represented as number of rows by number of columns.

#### Diseases

We limited our disease set to those diseases that have a known drug or a gene association (KEGG, RepoDB, DisGeNET, FDA, and DrugCentral). The diseases were mapped to the Unified Medical Language System (UMLS) concept unique identifiers (CUI).

In this study, we collected 5 different drug-related and one disease-related annotation profiles. We represent each annotation profile as a binary adjacency matrix. For 4008 drugs extracted from DrugBank [16], we extracted 2908 targets from DrugBank [16] and KEGG [17]; 881 chemical substructure profiles from PubChem [18]; 11744 side-effect annotations from SIDER [19] and OFFSides [20]; and 2124 MeSH (Medical Subject Heading) category profiles from DrugBank [16]. For the selected 2958 diseases, we downloaded their 7913 associated genes from KEGG [17] and DisGeNET (curated) [21]. Our drug-disease associations consists of 8984 indication pairs (4008 drugs and 2958 diseases) compiled from KEGG [17], FDA [22], DrugCentral [23], and RepoDB [24]. Each drug-related and disease-related annotation profile is considered as a *view* and we performed multi-label classification by classifying drug nodes to disease labels as associations using a multi-view learning approach.

### 2.2 Graph Neural Networks For Multi-views

In recent years, significant advances have been made in developing several variants of graph neural networks (GNN). The 1-dimensional Weisfeiler-Lehman test of isomorphism [25], also known as “naive vertex refinement”, is a powerful algorithm to test whether two graphs are isomorphic. It works by iteratively aggregating the labels of nodes and their neighborhood, and then performs hashing to the aggregated labels into unique labels. If at any iteration, the unique labels tend to be different for the two graphs, then the algorithm declares them as non-isomorphic graphs. The GNNs can also be viewed as a special case of 1-WL algorithm where neural networks are used to aggregate node neighborhoods [25], [26]. Compared to traditional machine learning models, GNNs have a greater capacity to identify interesting patterns from data. Given an adjacency matrix *A*, the graph convolutional network (GCN) in [8], a variant of GNN, performs semi-supervised graph learning by computing the transformation at each layer as

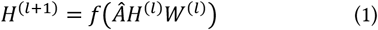

Where *Â* is obtained by “renormalization trick” given as 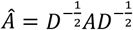. In simple terms, this is the preprocessing step before feeding the input *A* to the neural network. *H*^(*l*)^ is the hidden layer i.e., 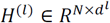. *d*^*l*^ is embedding size at each layer for *N* number of nodes, 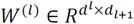, and *f*(.) is an activation function (Sigmoid or ReLU). This propagation mechanism emphasizes that each node will exchange first-order information in every layer followed by some non-linear transformation.

Several studies such as [9], [27], [28] extended the GCN framework to multi-view network. MedGCN [10], for instance, applied GCN on a multi-view network by learning node embedding for each heterogeneous node type by generalizing previous works neighborhood aggregation mechanism [29]–[31]. The GCN layer for multi-view network is defined as:

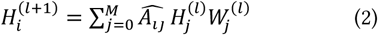

where 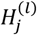 is hidden layer at layer *l* for node type *j, M* is the number of views, 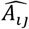 is the adjacency matrix for node type *i* and 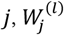 is a train-able weight matrix in layer *l*, and since the views are in bi-adjacency structure, random walk normalized Laplacian is applied i.e., *Â = D*^−1^*A*, is chosen as “re-normalization trick” for equation 2. Based on this generalized neural network approach for multi-view networks, we propose MGATRx consisting of two phases, the encoding and decoding phases.

### 2.3 MGATRx: A Multi-View Graph Attention Network

#### 2.3.1 Encoding

In GCN or multi-view GCNs, each node’s embedding is based on weighted sum of its neighbor embeddings. Previous studies from Velickovic et al. [11] and Thekumparampil et al. [12] used the attention mechanism for their aggregation process. Attention prioritizes neighbors with more relevant information to calculate embedding for each node. Thekumparampil et. al. proposed similarity-based attention where relevant neighbors that share similar representations are given more attention according to their contribution. Given two nodes *i* and *j*, the attention from node *j* to node *i* is calculated as

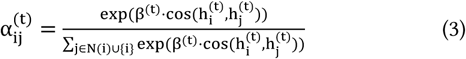

Where *h*_*i*_, *h*_*j*_ are the node embeddings of node *i* and *j*. In other words, attention (α_*ij*_) is obtained by multiplying a trainable scalar parameter *β* to the cosine-similarity of the embeddings followed by applying a SoftMax function on top of it for every layer *t*.

However, this model [12] was proposed for homogeneous graphs with a single-view. In the drug-disease association paradigm, the nodes are heterogeneous and have multiple views. Hence, the nodes should be able to attain relevant information from their heterogeneous neighborhood, filtering out any ‘useless’ information. In Figure 2, we illustrate a sample sub-graph of Simvastatin drug, prescribed to treat increased blood cholesterol levels (hypercholesterolemia). Given, valproic acid (an anti-epileptic) and liver carcinoma which also share the target HMGCR, one hypothesis is that simvastatin would exhibit promiscuity in treating new diseases while filtering irrelevant information. This hypothesis can be extended for each of the 4008 drugs. In other words, the attention mechanism is extended to multi-view networks as follows. Given *K* node-types (drugs, diseases, targets, mesh-categories, substructures and side-effects) in *M* bi-adjacency views, each node type feature matrix *X*_*i*_ is multiplied with trainable weight matrix W_*i*_ ∈ *R*^*N*×*d*^ to get an embedding (*H*_*i*_) of size *d*,

**Figure 1:**
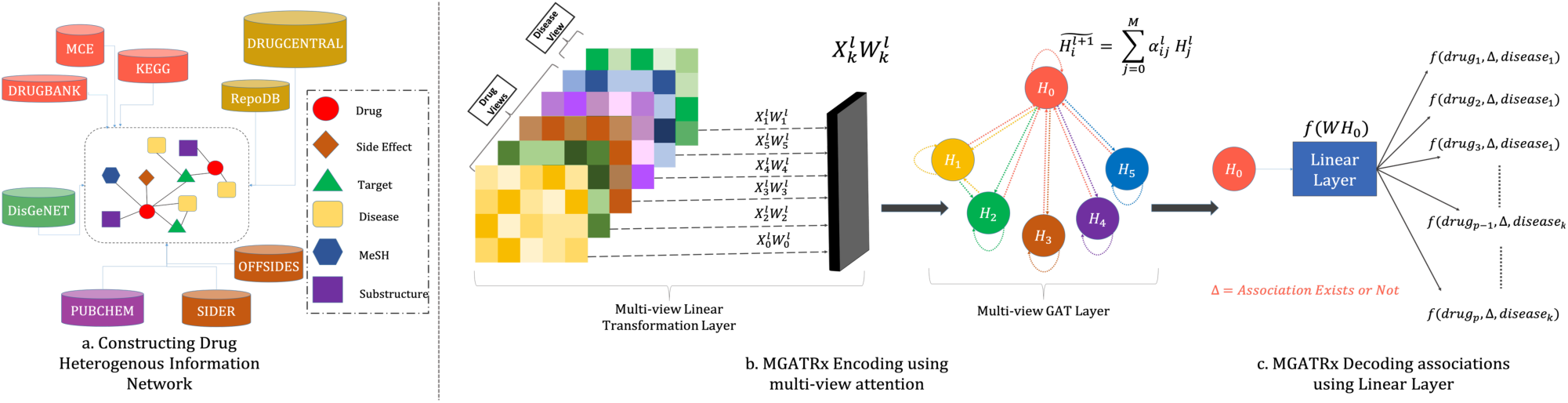
MGATRx architecture for predicting drug-disease associations. Panel a shows the drug-disease heterogeneous network construction step leveraging various data sources. Panel b represents our encoding algorithm which generates embeddings for multiple node types based on multi-view neighborhood information through attention. Panel c shows an overview of the decoding step wherein a linear layer is used as a decoder to decode the drug-disease relationships.

**Figure 2:**
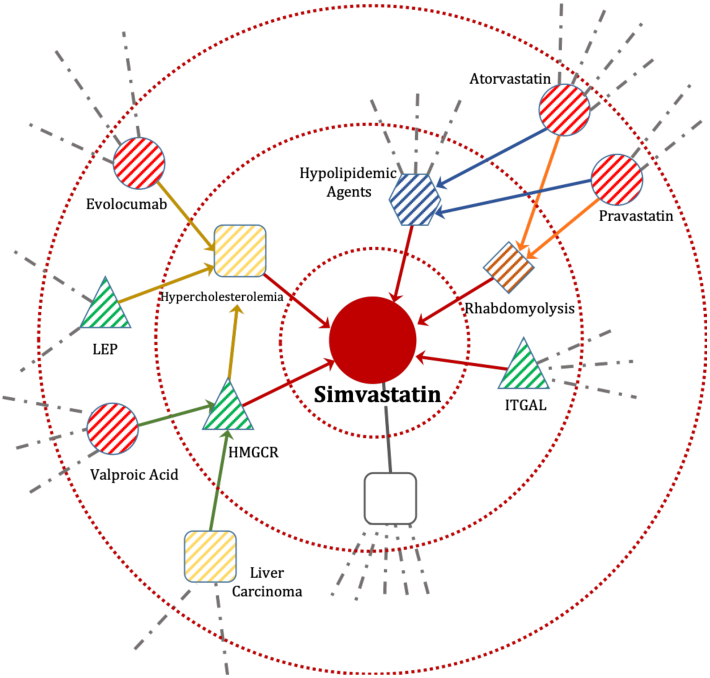
An example of MGATRx neighbor aggregation mechanism for the drug simvastatin. The circular nodes are drugs, squares are diseases, triangles are targets/genes, rhombuses are side-effects and hexagons are drug classes. The 1-hop and 2-hop heterogeneous neighbors of simvastatin drug are represented as dashed lines within the respective nodes. The incoming colored lines (with colors similar to the node they are directed to) represent the node learning its representations from the neighboring nodes. For instance, red colored edges towards simvastatin node represent that drug-associated representations are learnt from its neighbors (HMGCR, ITGAL, rhabdomyolysis, etc.).

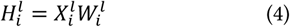

Since there are no external features for the *K* node types, we chose identity matrix as a node feature matrix. We apply attention (equation 3) and extend the multi-view sum pooling (equation 2) as multi-view graph attention. This way the attention mechanism over views learns which neighbors are more relevant and weighs their contributions accordingly. For node-type *i, j* ∈ *K*, each node learns its representation through inter (*i* ≠ *j*) and intra-attention (*i = j*) across the views as follows:

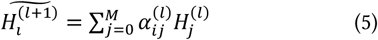

Here, we use softmax normalized attention *α*_*ij*_, as in equation 3 [12], which indicates the importance of nodes in view *j* to *i* i.e., 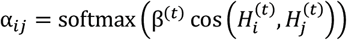. Similar to equation 3, 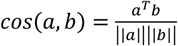 is the cosine similarity of the hidden states and *β*^(*t*)^ ∈ *R* is a trainable scalar parameter applied for *t* ∈ {1, …, *T*} propagation layers. The propagation layer (*t*), is different from embedding layer (*l*), learns to weight more relevant neighbors every time α_*ij*_ is computed. In single-view graph attention network [12], attention is computed for more than 1 propagation layer (*t* > 1). However, in our experiments we set the number of layers (*l* ∈ *L*) and attention propagation layers (*t* ∈ *T*) for each node type to *T =* 1 and *L =* 1. Finally, we applied scaled exponential linear unit (SELU) activation function for the attained embedding. Our choice of SELU is based on experimenting with various other activation functions provided in PyTorch.

#### 2.3.2 Decoding

In the decoding phase of MGATRx, we perform multi-label classification using a single linear layer i.e., *f*(*WH*_*i*_), where *W* is the weight matrix for reconstructing the adjacency matrix. Here, *f*(.) is sigmoid activation function which computes association score between the two node types i.e., 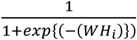. We tried DistMult, Inner Product and Bi-linear decoders for link-prediction between drug and disease but based on AUPR (Area Under Precision Curve) and AUROC (Area Under Receiver Operating Curve), the linear layer performed better than other decoders.

#### 2.3.3 Loss Function

For predicting drug-disease association, we calculate binary cross-entropy error (ℒ_ℬ𝒞ℰ_) of drug-disease association prediction and re-construction error of the remaining views using mean squared error (ℒ_ℳ𝒮ℰ_). The collective loss for MGATRx is given as

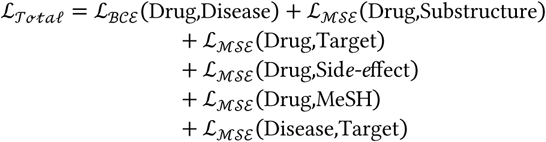

The binary cross entropy error ℒ_ℬ𝒞ℰ_ and mean-squared error ℒ_ℳ𝒮ℰ_ are defined as

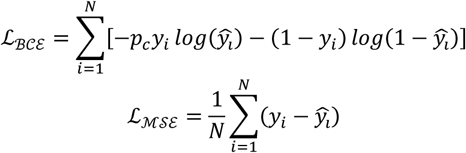

where *y*_*i*_ ∈ {0,1} is the ground truth, 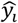 is the prediction for the *i*^*th*^ label, and *p*_*c*_ in the ℒ_ℬ𝒞ℰ_ are the weights for the positive labels to handle sparseness i.e., 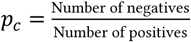

## 3 Experimental Setup

### 3.1 Hyperparameter settings for MGATRx

We implemented our MGATRx model in PyTorch. In our experiments, we adopted ten-fold cross-validation (10-CV) to test and compare performances of prediction models. We used stratified cross-validation from scikit [32] to distribute known and unknown associations in each fold. In each fold, we have training and testing sets. Further, from the training set, we used 15% of the data as validation set for hyper-parameter tuning. We constructed the prediction model based on known associations in the training set and predicted associations in the test set. We chose our best performing embedding size for each view as 512 after evaluating a set of dimension sizes [32, 64, 128, 256, 512]. In our experiments, we used Adam optimizer and performed hyper-parameter tuning with learning rate (η) set to 0.01 which was selected from a range of {0,1, 0.01, 0.001, 1e-4, 1e-5, 1e-6}, dropout set to 0.0 (no-dropout) which was selected from range of {0.0,0.1, 0.3, 0.5, 0.7, 0.9, 1.0} and weight-decay set to 0.0 which was selected from a range of {0.0,1e-3, 1e-3, 3e-3, 3e-5, 5e-3, 5e-5} parameters. During training, we used 2000 epochs in each fold where early stopping strategy is performed if AUPR does not increase for 50 successive epochs. For training, we used a workstation with Intel(R) Xeon(R) W-2133 CPU 3.60GHz CPU, 64 GB RAM and Nvidia Quadro RTX 8000 GPU.

### 3.2 Baseline Methods

For evaluation and comparison of our method, we used some of the state-of-the-art methods such as LRRSL, NeoDTI, and HNRD which are proposed for drug-repositioning problem. These baseline models are openly available in GitHub repositories. We did not use PREDICT [2] as a baseline because of the reproducibility issues (reported recently in OpenPREDICT case study [33]). To ensure fairness, we split the same size of training, validation, and test sets during 10-fold cross-validation.

**LRRSL** [5], proposed by Liang et. al, uses drug-related annotation profiles to identify drug-disease associations. In this study, as part of pre-processing step, they retrieve graph k-nearest neighbors (KNN) by calculating cosine similarity of the drug-annotation profiles. With the attained graph-knns, Laplacian is computed for each profile. The objective function is then defined as follows,

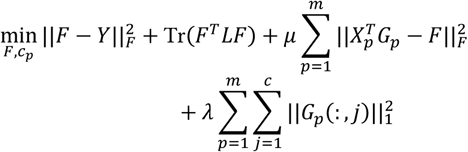

Where *X*_*p*_ represents the *p*^*th*^ view of drug annotation, *Y* is the drug-disease association matrix and *L* is the Laplacian matrix. *G*_*p*_ is the projected space where dimension is equal to the number of diseases to solve the drug-disease association scores in *F*. We used the default parameters mentioned by the work i.e., μ *=* 0.01, λ *=* 0.01, γ *=* 2 and *k =* 10.

**NeoDTI** [34] by Wan et.al predicts drug-target association. NeoDTI learns node-level embedding by aggregation technique with respect to node-type and predicts the confidence score between a drug and a protein as

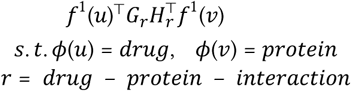

where *G*_*r*_, *H*_*r*_ are learned edge-type specific projection matrices, ϕ(*u*) is drug node type, ϕ(*v*) is protein node type and *r* is the relation. Although, NeoDTI is proposed for drug-target prediction, the model can be extendable to predict drug-disease associations. Note that DTINet [35], a study preceding NeoDTI, uses similar reconstruction strategy by unifying multiple networks to predict drug-target associations. However, in the performance comparison with NeoDTI, DTINet was reported to be inferior to NeoDTI. Hence, we did not consider DTINet as a baseline in the current study.

**HNRD** [7] proposed by Wang et. al utilizes NeoDTI affinity-based approach of neighborhood aggregation to predict drug-disease associations based on drug-drug and disease-disease similarity networks.

**MedGCN** [10] from Mao et. al uses multi-view graph convolutional network for medication recommendation problem as discussed in section 2.2. The multi-view GCN aggregates neighbors for each node-type as 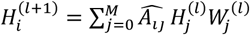.

### 3.3 Evaluation Metrics

We used the traditional evaluation metrics such as area under the precision-recall curve (AUPR) and area under receiver operating characteristic curve (AUROC) to quantitatively evaluate and compare our method with the existing baseline methods. We identified optimal threshold based on elbow method of AUPR curve and computed F1-score. To evaluate the top-k predictions, we report average precision at K. Higher values in Table 2 indicate better performance.

**Table 2:** Performance comparison using 10-fold cross-validation setting.

| Methods | AUPR | AUC | F1 | AP@1 | AP@5 | AP@50 |
| --- | --- | --- | --- | --- | --- | --- |
| LRSSL | 0.212 | 0.928 | 0.283 | 0.779 | 0.660 | 0.637 |
| NeoDTI | 0.385 | 0.902 | 0.448 | 0.609 | 0.505 | 0.482 |
| HNRD | 0.421 | 0.933 | 0.467 | 0.552 | 0.444 | 0.428 |
| MedGCN | 0.471 | 0.925 | 0.511 | 0.837 | 0.750 | 0.730 |
| **MGATRx (Ours)** | **0.568** | **0.944** | **0.586** | **0.858** | **0.791** | **0.779** |

## 4 Results

### 4.1 Performance Evaluation and Comparison with Other State-of-art Approaches

Evaluation and comparison of MGATRx with current state-of-art methods using the hyper-parameters as provided in the literature showed superior performance of MGATRx (Table 2). LRSSL showed a notable AUC performance compared to other baseline methods, but in terms of AUPR and F1, the scores were inferior. Notably graph neural network-based methods (NeoDTI, HNRD, MedGCN, and MGATRx) showed better AUPR and AUC performance. It is, however, surprising to notice that similarity-based method of NeoDTI i.e., HNRD performed better than the original. The performance of MedGCN is inferior when compared to MGATRx suggesting that attention helps in exploiting higher-order neighbors in multi-view networks.

To test the argument that our proposed attention model is robust than convolution-based method, we removed a fraction of edges and compared the performance. In Figure 3, we show the robustness of MGATRx and MedGCN, by removing 15%, 30%, 45%, 60% and 75% of the edges. Note that the removed edges were used as validation set during training and performed early stopping if the model performance over validation set did not improve for 50 epochs as discussed in section 3.1. Our proposed algorithm performed substantially better than MedGCN in robustness test, despite removing edges from the training set and evaluated with test set (Figure 3). Besides, MGATRx also has an additional advantage of attention interpretation over MedGCN (discussed in section 4.3). Overall, MGATRx shows a superior performance of at least 8-35% in terms of AUPR scores when compared with the baseline models.

**Figure 3:**
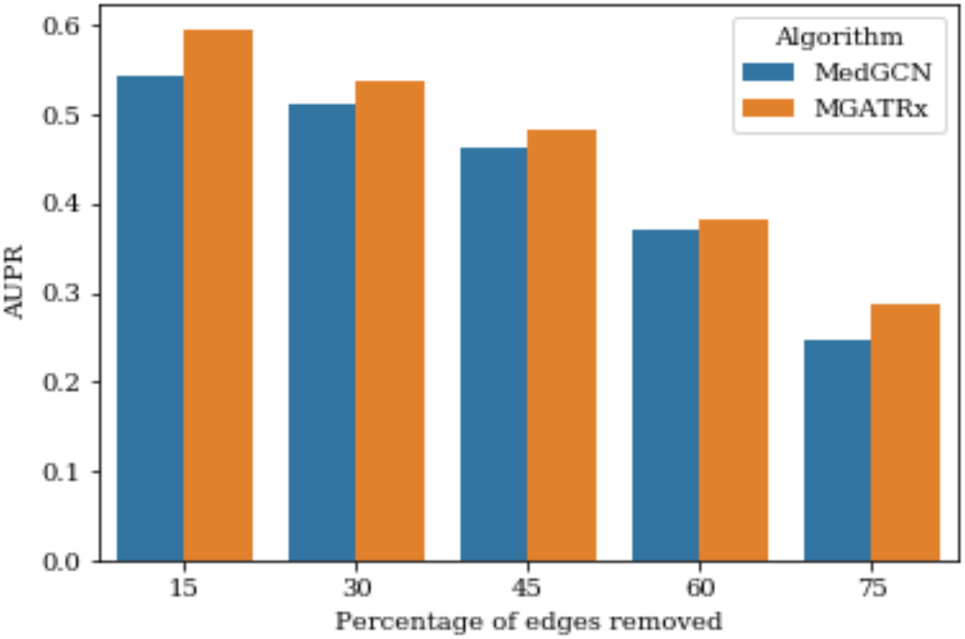
Results from robustness analysis performed by removing percentage of edges.

### 4.2 Hyper-parameter Influence and Analysis

In this section, we will analyze the hyper-parameters influencing the prediction of MGATRx namely, learning rate (lr), dropout, weight decay and embedding size. For learning rate, dropout and weight decay parameters, the plots in Figure 4 (a), (b) and (c) are first-fold validation AUPR scores. This demonstrates the convergence of the solution at every epoch step. In case of embedding size, we evaluated the 10-fold AUPR scores for each embedding size and represented as box plot.

**Figure 4:**
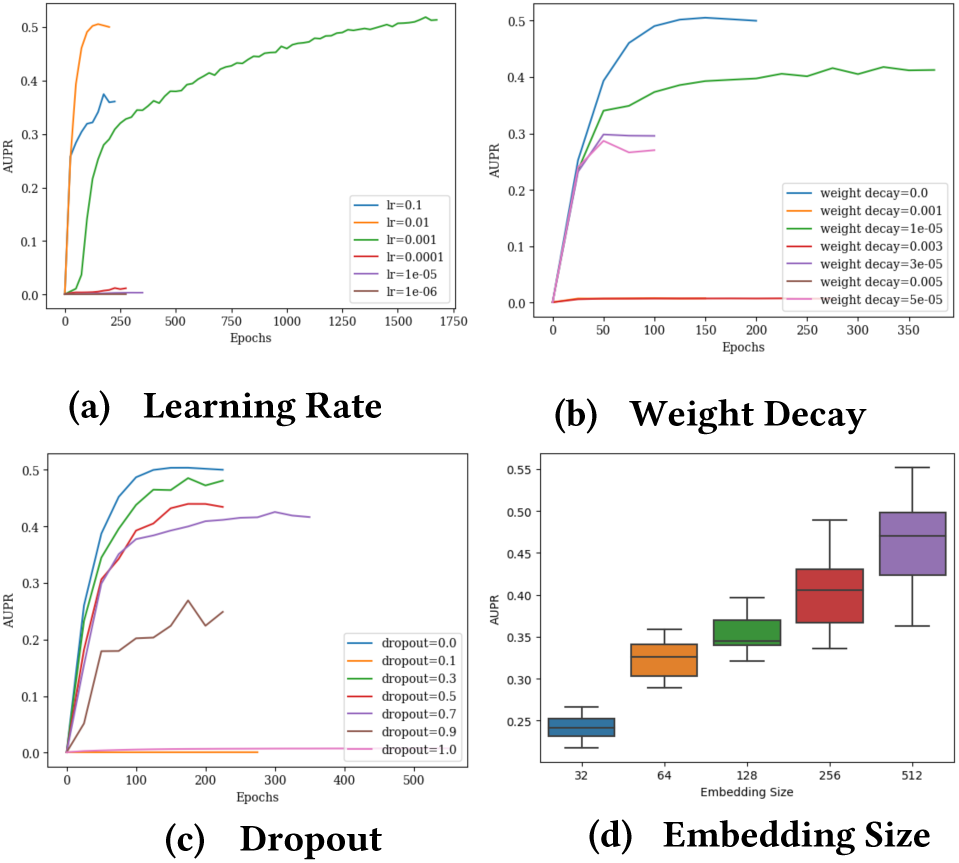
Hyper-parameter tuning and analysis for MGATRx.

#### Learning Rate

As shown in Figure 4 (a), MGATRx can converge solution with an optimal learning rate at η *=* 0.01. The algorithm can achieve performance with larger learning rate. However, as the learning rate is decreasing, the performance decreased gradually. This variance in the performance could be attributed to Adam optimizer’s momentum and adaptive learning rate tricks.

#### Weight Decay

Like dropout, weight decay is tuned based on a search space of seven parameters. We observed that with no weight-decay the network can perform well.

#### Dropout

For dropout we used a range of dropout values and found that the dropout at its lowest value set to 0.1 performed better than others. Due to the sparsity in the graphs, increase in the dropout affects the performance of the model as shown in Figure 4 (c).

#### Embedding Size

In our experiments, we observed that keeping the rest of parameters constant, the embedding size influenced the AUPR performance. With embedding size set to 512 the performance is relatively superior when compared with rest of the models.

### 4.3 Interpreting Attention

To visualize the impact of attention on the model, we selected immunological disorders such as asthma, psoriasis, Crohn’s disease, and chronic ulcerative colitis. We selected top 5 predicted drugs for each of these diseases based on drug-disease attention weights. Next, we filtered 15 targets which are shown to have maximum attention from these drugs and diseases. In Figure 5, we show the drug-target and disease-target attention heatmap. The highlighted values in the heatmap show how much attention a target gene is contributing for constructing drug and disease representations, which further aids in predicting association between a drug and a disease. For example, we selected the highlighted attention targets showed in the heatmap for “Chronic Ulcerative Colitis” and then used ToppGene Suite [36] to identify functional enrichments for biological processes. We found that these genes are enriched (p value < 0.05 FDR) for inflammatory response (GO:0002437) and regulation of immune process (GO:0050776). This probably explains the presence of immune suppressive agents (such as Natalizumab, Adalimumab, Infliximab, Risankizumab, Sarilumab, and Certolizumab Pegol) among the top hits for these immunological disorders. Similar enrichments were observed for Crohn’s disease. Certolizumab pegol, an approved drug for Crohn’s disease, has been reported to be also efficacious for treating ulcerative colitis [37], [38]. The gene TNF appears to be an active contributor of attention to both diseases. Likewise, in case of asthma, IL5 is observed to be one of the genes contributing attention factor. Mepolizumab and Reslizumab are approved drugs for asthma and therefore it is not surprising to see IL5 contributing maximum attention to the drugs. It is to be noted that the current analysis is primarily based on targets and their contribution to drugs and diseases. However, there are other node-types such as side-effects, chemical structure, and MeSH categories. Since, targets are the common space for both the drugs and disease, we used them for this qualitative elucidation.

**Figure 5:**
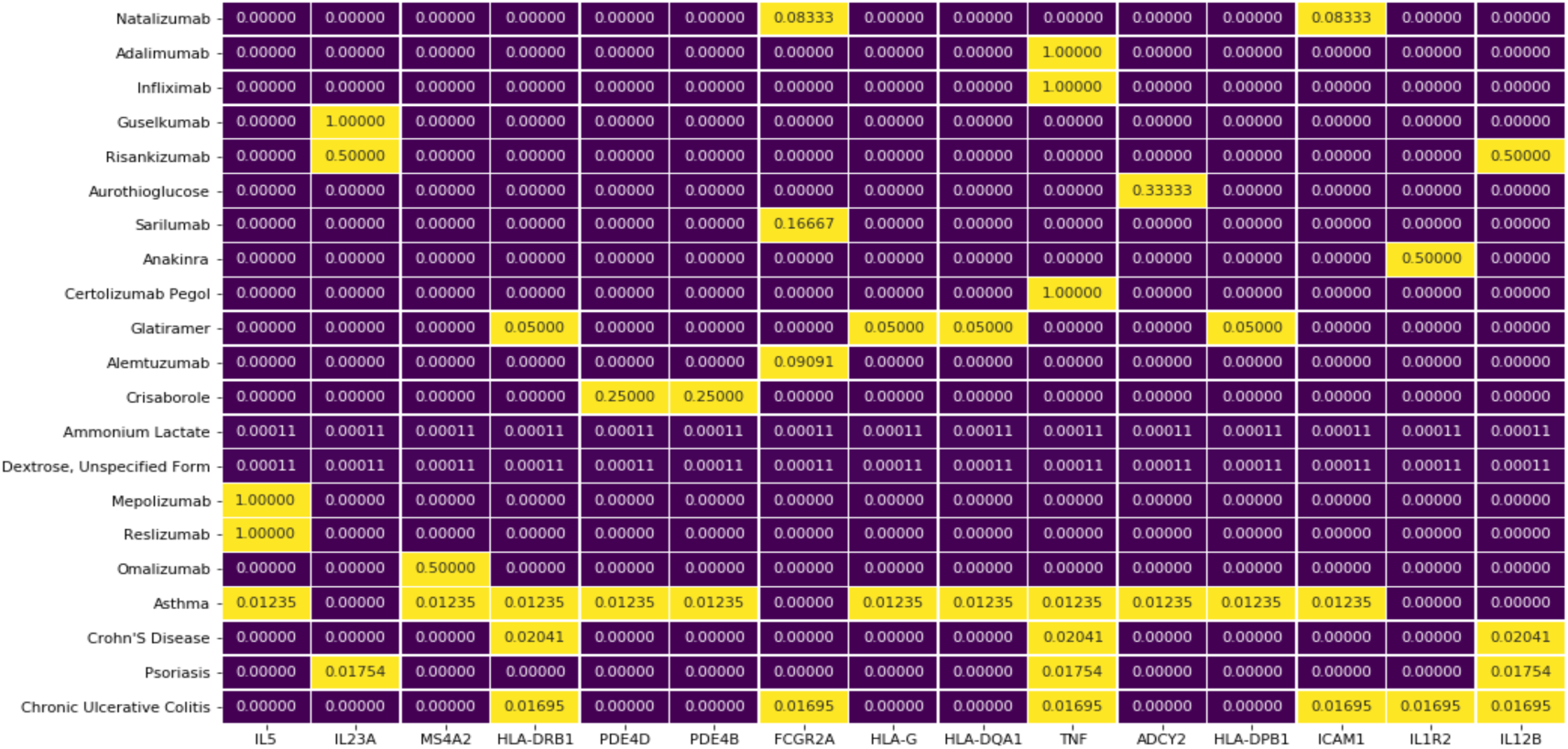
Drug and Disease attention weight visualization based on targets. The attention scores are in the range of {0,1}

### 4.4 T-SNE Visualization

We extracted the drug representation in the decoding layer and applied t-SNE (t-Distributed Stochastic Neighbor Embedding) to further identify hidden patterns. In Figure 6, we show the clusters of drugs in 2-dimensional Euclidean space. To color code the drugs, we used Anatomical Therapeutic Chemical (ATC) codes, which is a drug classification system adopted by World Health Organization (WHO). There are 14 ATC codes and each code has 4 sub-levels. We extracted ATC codes from DrugBank and used the first level of the code. We assign color codes to each drug based on ATC Code and observe few clusters of drugs such as nervous system drugs in blue, cardiovascular drugs in red, anti-neoplastic drugs in teal and anti-infectives in grey. The visualization also reveals potential promiscuity patterns of drugs such as alimentary tract, anti-infectives and anti-parasitic related drugs. These overlaps also suggest drug repositioning potential (drugs are like other ATC class).

**Figure 6:**
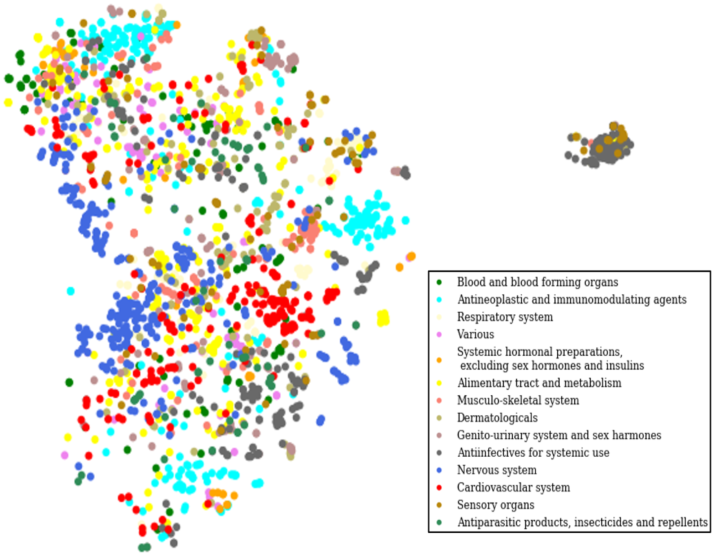
2-Dimensional T-SNE visualization of drug representations. There are 14 ATC codes and each drug is mapped according to its ATC code.

### 4.5 Novel indications and drug repositioning candidates

In this section, we briefly discuss some of the novel indication predictions of MGATRx. We selected 12 drugs with their original indication and MGATRx-predicted indication along with literature evidence for on-going investigations (Table 3).

**Table 3:**
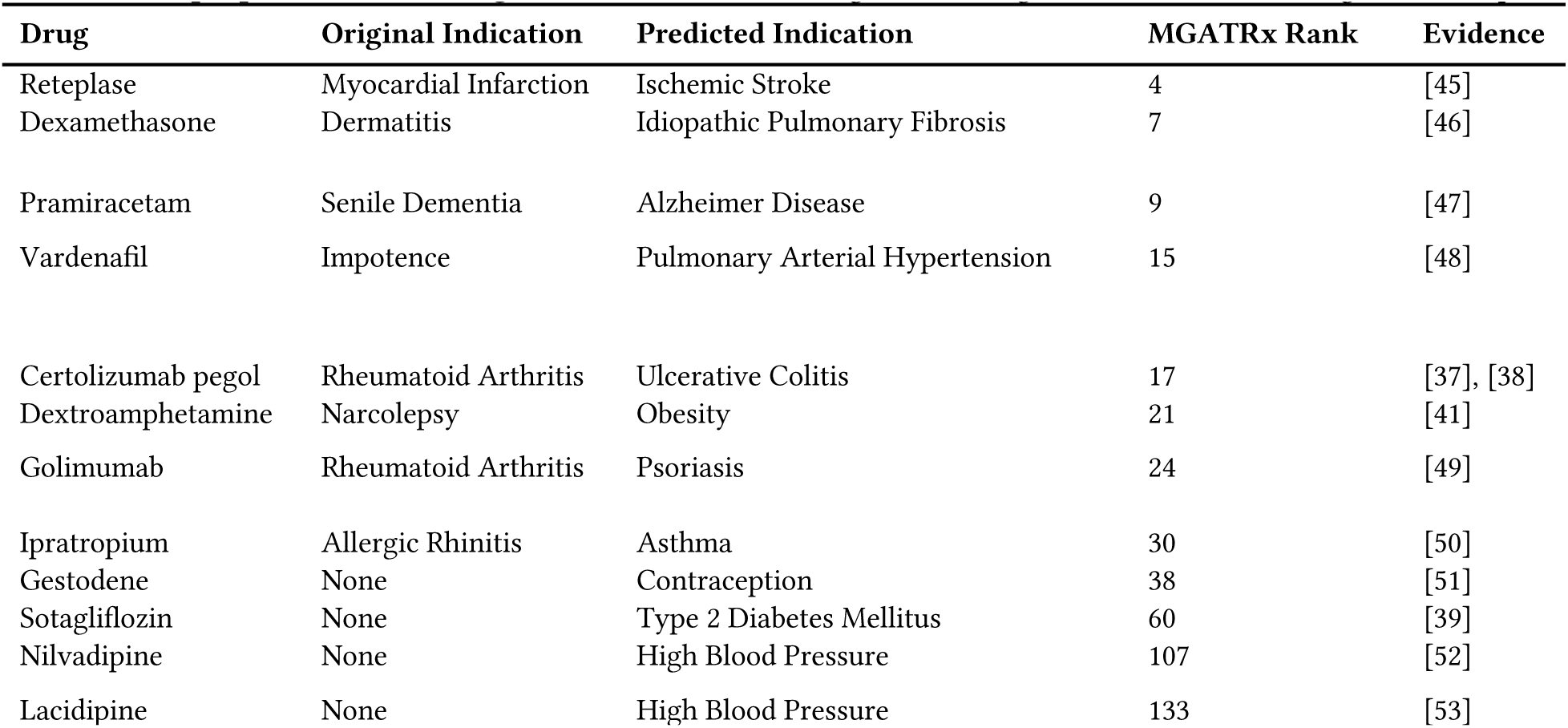
Example predictions for drug-disease associations. Drugs with no original indication are investigational compounds

| Drug | Original Indication | Predicted Indication | MGATRx Rank | Evidence |
| --- | --- | --- | --- | --- |
| Reteplase | Myocardial Infarction | Ischemic Stroke | 4 | [45] |
| Dexamethasone | Dermatitis | Idiopathic Pulmonary Fibrosis | 7 | [46] |
| Pramiracetam | Senile Dementia | Alzheimer Disease | 9 | [47] |
| Vardenafil | Impotence | Pulmonary Arterial Hypertension | 15 | [48] |
| Certolizumab pegol | Rheumatoid Arthritis | Ulcerative Colitis | 17 | [37], [38] |
| Dextroamphetamine | Narcolepsy | Obesity | 21 | [41] |
| Golimumab | Rheumatoid Arthritis | Psoriasis | 24 | [49] |
| Ipratropium | Allergic Rhinitis | Asthma | 30 | [50] |
| Gestodene | None | Contraception | 38 | [51] |
| Sotagliflozin | None | Type 2 Diabetes Mellitus | 60 | [39] |
| Nilvadipine | None | High Blood Pressure | 107 | [52] |
| Lacidipine | None | High Blood Pressure | 133 | [53] |

Among the MGATRx predicted novel indications is type 2 diabetes (T2D) for the drug Sotagliflozin. Interestingly, this drug is currently approved in Europe [39] but not yet by the FDA. Similarly, calcium channel blockers (CCB) such as Nilvadipine and Lacidipine are predicted for high blood pressure. Most CCBs are known to have therapeutic efficacy for hypertension [40]. Among other examples, Dextroamphetamine (DAH) is a well-known drug for treating narcolepsy and attention deficit hyperactivity disorders. In our predictions, we found that DAH is predicted to treat obesity. Recent studies report beneficial effects of DAH in treating hypothalamic obesity in children and adolescents [41], [42]. Similarly, Certolizumab pegol (CMZ) is predicted as a potential repositioning candidate for ulcerative colitis. CMZ is a known anti-TNF agent and in our model, attention towards TNF target was observed (described in earlier sections). CMZ is currently under active investigation in phase II clinical trial [43].

The attention contributed by a known target or an off-target may not necessarily be always a potential candidate for repositioning. For example, some of the anti-TNF drugs such as Infliximab and Adalimumab are reported to be associated with aggravating multiple sclerosis in patients [44]. However, MGATRx ranks them among the top predictions. This limitation, we believe, can be potentially addressed in future work by incorporating the directionality (e.g., drugs treating a phenotype vs. drugs causing a side-effect or disease-causing a phenotype, etc.) to the edges while representation in the model. The goal of MGATRx is to generate testable hypotheses for drug repositioning for potential experimental validation,

## 5 Conclusion

In-silico drug repositioning is a critical component of all drug discovery pipelines. Most of the previously reported similarity-based approaches for predicting drug-disease associations assume that nodes sharing similar annotations could have similar associations. Most of these approaches further use relatively older versions of drug-disease data which had limited coverage apart from being not up-to-date. To address this, we systematically curated annotations from various sources and constructed a multi-view heterogeneous network. Leveraging this relatively current drug and disease annotations, we built MGATRx, a novel approach to predict and identify drug repositioning candidates. MGATRx is a multi-view graph attention drug repositioning framework which selectively aggregates relevant information from its neighbors to learn node representations. Our comparative analysis with four current state-of-art methods shows a substantial improvement in prediction performance. Besides, we also visualized the learned attention for select drugs and disease groups to enable understanding of the molecular basis for the drug-disease predictions. Several of our predicted drug-disease indication pairs overlap with drug indications that are either currently in clinical trials or are supported by literature references, demonstrating the overall translational utility of MGATRx. Apart from striving to update the disease-drug annotations regularly, as part of future plans, we seek to expand the MGATRx framework to identify drug combinations (e.g., synergistic or non-synergistic drug combinations) or drug-induced adverse events, or polypharmacy-induced adverse events. Although the multi-view graph attention achieved superior results for drug repositioning, it would be interesting to consider other drugs, diseases, and gene annotations such as drug and/or disease-related transcriptomics data, target variant, protein interactions, pathways, and drug dosage-relevant information as additional views. Including these additional views, we hypothesize, may not only enhance the MGATRx repositioning performance but also enable our understanding or formulating hypotheses of drug-disease mechanistic relationships. Lastly, but most importantly, in support of reproducible research endeavor, we make all the MGATRx data and source code publicly available^1^.

## ACKNOWLEDGEMENTS

This work was supported, in part, by NIH NCATS grant 1UG3TR002612 to AJ.

## Footnotes

1 https://github.com/yellajaswanth/MGATRx

